# Composition of saliva metabolome is significantly associated with SARS-CoV2 infection and with severity of COVID-19 disease

**DOI:** 10.1101/2024.10.24.620087

**Authors:** Violeta Larios-Serrato, Natalia Vázquez-Manjarrez, Osbaldo Resendis-Antonio, Nora Rios-Sarabia, Beatriz Meza-Marquez, Oliver Fiehn, Javier Torres

**Affiliations:** Departamento de Bioquímica, Escuela Nacional de Ciencias Biológicas, Instituto Politécnico Nacional, Mexico City, México; Unidad de Metabolómica, Dirección de Nutrición, Instituto de Ciencias Médicas y Nutrición Salvador Zubirán, Ciudad de Mexico 14080, Mexico City, Mexico; Human Systems Biology Laboratory, Instituto Nacional de Medicina Genomica & Coordinación de la Investigación Científca-Red de Apoyo a la Investigación, UNAM, Mexico City, Mexico; Unidad de Investigación Médica en Enfermedades Infecciosas, UMAE Pediatria, Centro Médico Nacional SXXI, IMSS, México; Universidad Autónoma de Baja California Sur, México; Centro de Investigaciones Biológicas del Noroeste SC, México; UC Davis West Coast Metabolomics Center, Davis, CA, USA

**Author notes:** Corresponding author: Javier Torres.

**Keywords:** metabolome, COVID-19, severity, patient

## Abstract

**Background:** The metabolome of COVID-19 patients has been studied sparsely, with most research focusing on a limited number of plasma metabolites or small cohorts. This is the first study to test saliva metabolites in COVID-19 patients in a comprehensive way, revealing significant changes linked to disease severity and highlighting saliva is potential as a non-invasive diagnostic tool.

**Methods:** We included 30 asymptomatic subjects with no prior COVID-19 infection or vaccination, 102 patients with mild SARS-CoV-2 infection, and 61 hospitalized patients with confirmed SARS-CoV-2 status. Saliva samples were analyzed using hydrophilic interaction liquid chromatography-mass spectrometry (HILIC-MS/MS) in positive and negative ionization modes.

**Results:** Significant changes in metabolites were identified in COVID-19 patients, with distinct patterns based on disease severity. Healthy individuals exhibited a well-regulated bacterial network, while severe cases showed disordered microbial networks. Elevated dipeptides such as Val-Glu and Met-Gln in moderate cases suggest specific protease activity related to SARS-CoV-2. Increased acetylated amino acids like N-Acetylserine and N-Acetylhistidine indicate potential biomarkers for stress and disease severity. Bacterial metabolites, including muramic acid and indole-3-carboxaldehyde, were higher in mild-moderate cases, indicating oral microbiota changes. In severe cases, polyamines and organ damage-related metabolites, such as N-acetylspermine and 3-methylcytidine, were significantly increased. Interestingly, metabolites reduced in moderate cases were elevated in severe cases.

**Conclusions:** Saliva metabolomics offers insight into disease progression and potential biomarkers for COVID-19.

## Introduction

Four years have passed since SARS-CoV2 struck humankind. As of June 2024, 776 million people have been infected and almost 7 million subjects died. SARS-CoV2 has successfully adapted to its new host, with around 129,000 new cases per day as per June 2024, many of them vaccinated via multiple rounds of vaccines. For the first time, science has exhaustively registered the evolutive process of viral adaptation to humans, and the genome of millions of SARS-CoV2 variants has been sequenced (Harari et al. 2024). Thus, the emergence of highly diverse variants has been captured almost in real-time, offering the opportunity to identify specific variants of concern and to implement optimal public health measures to reduce their spread. The extent of analysis of the emerging variants may even allow predictions on mutants, particularly in the receptor binding domain, that will be circulating in particular populations.

With over 7 million deaths it is clear that early SARS-CoV2 episodes were fierce, with a high mortality rate. Our innate response was not trained to fight SARS-CoV2. It seems that an unregulated, explosive response to infections was an important cause for severe and even fatal cases, i.e. an uncontrolled production of IFN that led to severe disease progression by interfering with lung epithelial regeneration (Sievers et al. 2024). Currently, a human/SARS-CoV2 co-adaption process and vaccination programs have led to a reduced mortality rate and a more trained innate and adaptive response. A modulation of the inflammatory response is required to fight back the infection and reduce the damage to the patient (Tang et al. 2020).

Learning about the nature of the interaction human/SARS-CoV2 during the early periods of this encounter offers important information on the response of a naïve population to infection with an unknown virus of zoonotic origin, a threat that is forever latent (Mollentze and Streicker 2020). A key element in this scenario is the microenvironment at the site of infection, defined not only by the virus and the epithelial cells but also by the immune cells, secreted cytokines and antiviral molecules, which collectively contribute to the complex etiology of the disease. The upper respiratory tract is a niche with a rich microbial community in humans, second in diversity only after the gut microbiota (Dewhirst et al. 2010). This microbial community may modulate infectivity, transmission, and even severity of viral infections (Tsang et al. 2020). The microbiota presents a key element to define the composition of the metabolome in the respiratory tract microenvironment, either by directly producing specific metabolites or by influencing the metabolism of the epithelial and immune cells in the niche. Importantly, there are scarce reports on the metabolome of the respiratory tract in COVID-19 patients. We here set out to reveal the role of these metabolites in the outcome of the infection.

We recently reported major changes in the microbiota of the saliva of COVID-19 patients (Larios Serrato et al. 2023) and found that diversity, composition, and networking of the microbial communities differed according to the severity of the disease. The metabolic activity of these communities was inferred with Picrust2 software, and major differences were predicted, particularly in the metabolism of amino acids, fatty acids, and antibiotics. Major changes in metabolic pathways of bacterial communities would lead to pronounced changes in the microenvironment where the newcomer SARS-CoV2 is trying to colonize. A detailed characterization of the metabolome in the mouth should provide information to define the “culture medium” where the viral infection is established and give clues to unravel the pathogenesis and severity of the disease (Moreno et al. 2024).

In this work we aimed to thoroughly analyze the metabolome in saliva of patients that acquired SARS-CoV2 infection during the early phase of the pandemic, before the advent of the vaccine and without previous infection. In addition, we aimed to elucidate possible differences between cases with diverse severity of the disease. Results showed that SARS-CoV2 infection was associated with markedly significant changes in the composition of the metabolome, indicative of an explosive and intense response to the infection. Major changes were also observed in association with the severity of the disease.

## Material and Methods

### Patients studied

Patients were recruited between June 2020 to January 2021, before any vaccination program was in place and during a period of uncertain epidemiologic and clinical scenarios with strict containment measures. Therefore, patient recruitment had to be conducted by the attending health personnel of the Hospital General de Zona con Unidad Medicina Familiar No. 8 in Mexico City. Patients with suspicious symptoms before a confirmed diagnosis participated in the study. Those who consented were asked to sign an informed consent letter. The study was approved by the IRB Comision Nacional de Investigación, Instituto Mexicano del Seguro Social, México (registry number R-2020-785-053).

Individuals without symptoms and no previous COVID-19 infection were included in the asymptomatic cases group (AC). Smokers and those with any chronic disease were excluded. A group of mild cases included ambulatory patients with mild respiratory symptoms (fever, cough, headache, odynophagia, myalgias) presenting for COVID-19 diagnosis, and recruited before any prescribed treatment. After testing for SARS-CoV-2 (COBAS 6800, Merck Mexico, Mexico City), they were classified as ambulatory (mild cases) positive (AP) or negative (AN) for the infection. Patients who during follow up required hospitalization because they developed severe symptoms were excluded from this group and included in the hospitalized group. The group of severe patients was cases that required hospitalization because of severe symptoms (HP group), with oxygen saturation below 92% and the presence of risk comorbidities including hypertension, diabetes, morbid obesity, immunocompromised, cardiovascular or neurological diseases, chronic renal failure, tuberculosis or neoplasia. Patients were usually hospitalized within the first seven days after symptoms started and followed until discharged because of recovery, improvement, or death. A summary of the type of classification and numerical frequency in each case is shown in **Table 1**.

**Table 1.**
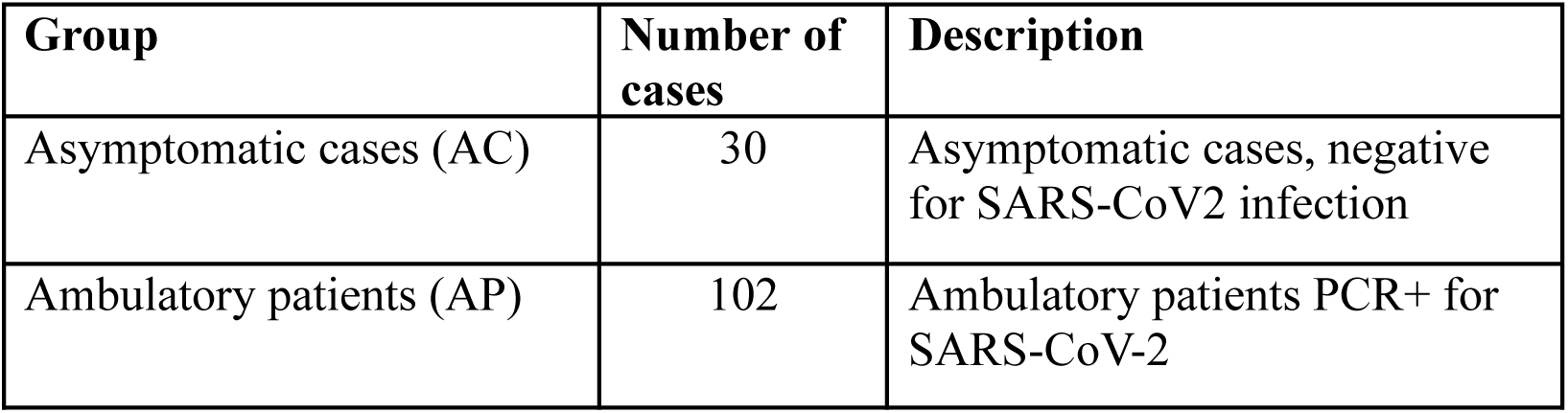
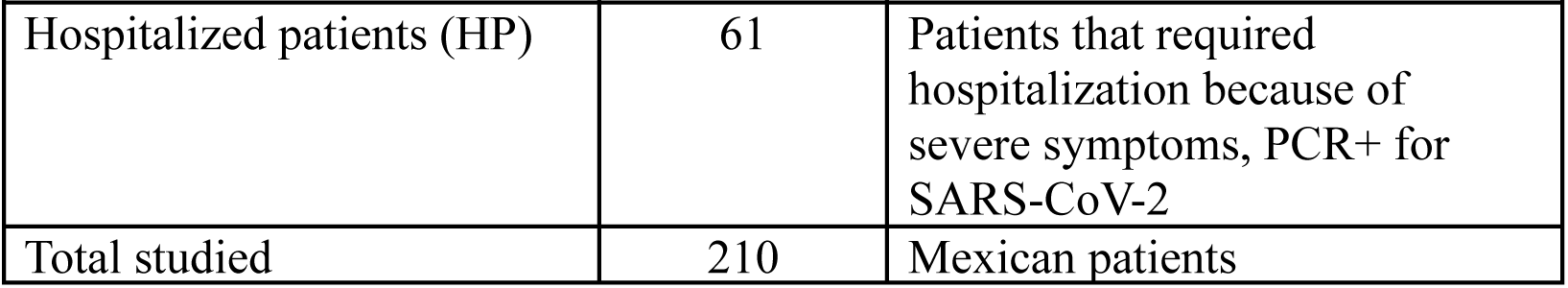
General characteristics of the clinical groups studied.

### Saliva samples

Patients were asked to thoroughly wash their mouths with 10 mL of saline solution (0.85% NaCl) and spit back into a 50 mL plastic tube. Samples were immediately transported to a central laboratory for SARS-CoV-2 diagnosis. On arrival, samples were inactivated by heating at 65 °C for 30 min. After the diagnosis, samples were sent to our laboratory following international safety regulations. Once received in our lab, samples were frozen at −70 °C until studied. The sample preparation procedure was performed according to standard protocols for untargeted metabolomics. After extraction, the samples were analyzed using hydrophilic interaction liquid chromatography-mass spectrometry (HILIC-MS/MS) in positive and negative ionization modes to generate the metabolomic data.

### Data Preprocessing and Quality Control

Data were acquired on a Sciex TripleTOF 6600 high resolution mass spectrometer, using a Waters BEH Amide 2.1 x 50 mm column with 1.7 um particles and an acetonitrile/wáter gradient (Blaženović et al. 2019). Raw LC-MS files were processed by MS-DIAL 4.0 software (Tsugawa et al. 2015). A total of 3,452 features were obtained in positive (2,218) and negative (1,234) mode. Ion intensities were then normalized by Systematic Error Removal by Random Forest (SERFF, (Fan et al. 2019)). Compounds were annotated by matching accurate mass and MS/MS data against NIST17 and MassBank of North America libraries. Principal Component Analysis (PCA) was as quality control to assess the overall sample distribution and detect any outliers or batch effects (**supplemental figure 1**).

### Statistical Analysis, Variable Selection, and Performance Assessment

The statistical analysis began with the selection of main ions pertinent to the biological question using Partial Least Squares Discriminant Analysis (PLS-DA) using MATLAB R2023b and PLS_toolbox 9.1. Only annotated ions were retained for further analysis, resulting in a size 308 x 905 data matrix. The data matrix was divided into a calibration set and a test set, with 10% of the samples assigned to the test set. PLS-DA was performed on the data matrix to discriminate between the groups, using Pareto-scaled data with 100 iterations and leave-one-out cross-validation for the inner loop to optimize the model’s parameters. The groups analyzed included: AC vs. AP, AC vs. HP and AP vs. HP (see **Table 1**). The selected features from PLS-DA were then applied to the test set to evaluate their discriminative performance. Performance metrics included the Area Under the Receiver Operating Characteristic Curve (AUROC) and the misclassification error. Variable selection was conducted using the selectivity ratio and Variable Importance in Projection (VIP) values. Features identified as highly discriminant in at least 80% of the models were considered relevant to the biological question.

Differences in metabolites between experimental groups were further analyzed to identify compounds that significantly distinguished each group (marker metabolites), R 4.4.0 packages and Rstudio 2024 were utilized. One-way Analysis of Variance (ANOVA) was used to explore the data with multigroups, and the False Discovery Rate (FDR) was calculated to identify important compounds among all comparisons (p < 0.05). Violin plots were built with the ggplot2 v3.5.1 package (Wickham 2016). The geometric mean for each metabolite across all samples was calculated using DESeq2 v1.44.0 (Love, Huber, and Anders 2014) and normalized using the total sum scale (TSS). To analyze the differences in the abundance of metabolites between groups, the fold change (FC) was calculated, and p-values were obtained using the Wald test. These values were corrected for multiple testing using the Benjamini and Hochberg method by default. Volcano plots were created with EnhancedVolcano v1.12.0.

### Data availability

Database of metabolome is described in Supplemental **Table 1**, showing data such as run information, intensities and internal standards.

## Results

### The composition of saliva metabolites clearly distinguished the clinical groups

We first determined if the composition of saliva metabolites could distinguish each clinical group. Interestingly, PCA analysis resulted in a clear separation of the three groups: the asymptomatic (AC), the moderate (AP), and the severe cases (HP) (**Figure 1**). The analyses show that component 1 explained 34.8% of the variance and component 2 the 16.2%, whereas components 3, 4 and 5 still had an important contribution (**Figure 1b)**. To identify the metabolites that significantly contributed to the observed separation of the groups, we performed a PLS-Da analysis in a pairwise comparison between groups (**Figure 2a**). Results showed that 35 metabolites distinguished AC from AP, 33 AC from HP, and 37 AP from HP, and a search in the literature revealed that the source of these metabolites might be human, environmental or microbial (**Table 2**). Furthermore, the predictive ability of these markers was tested by estimating the ROC value and all three pair comparisons resulted in excellent predicted ROC and response values (**Figure 2b**). **Supplemental Table 2** depicts the differences in intensity of the selected metabolites by group.

**Figure 1.**
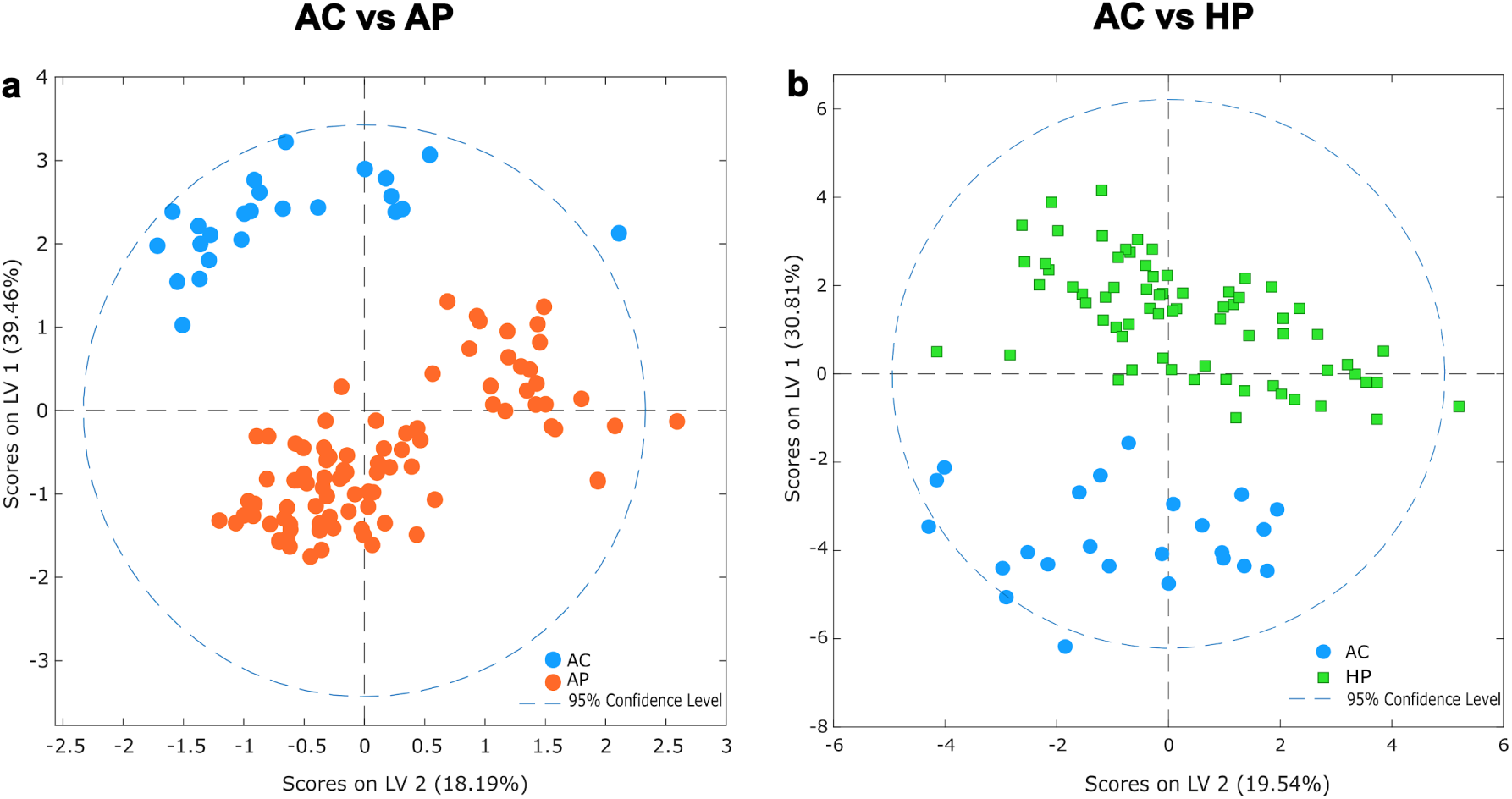
Score plots. Pairwise score plots between components of unsupervised method, component 1 shows the major separation between the groups AC, AP and HP.

**Figure 2.**
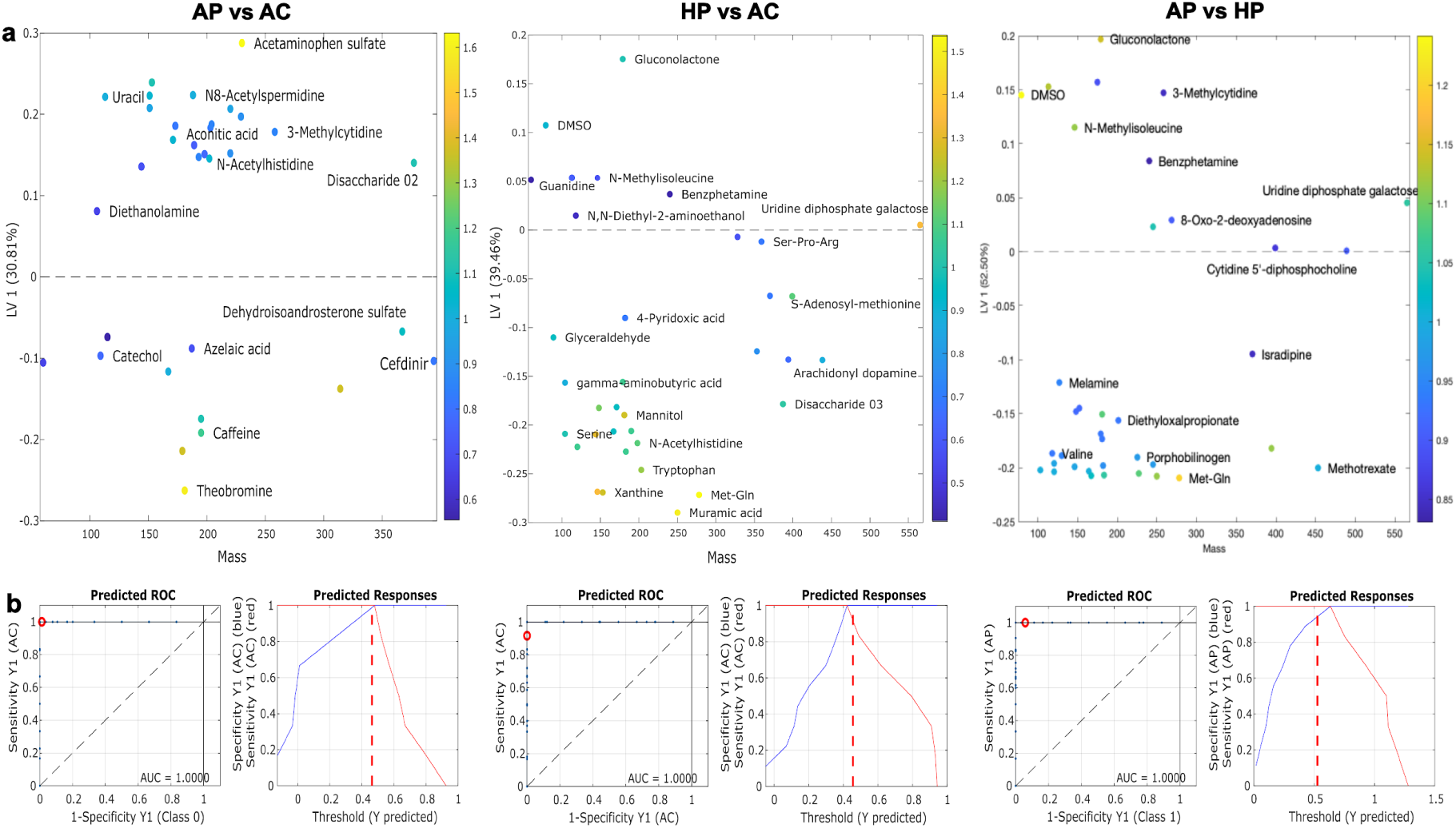
Metabolites with significant differences between the clinics groups using a Very Important Metabolites (VIP) analysis. (a) dot plots for each comparison, showing mass on x-axis and LV1 on y-axis. Dashed lines reflect threshold, LV ≥ .0 (abundance in reference group) and LV ≤ 0 (abundance in second group); AC vs AP, comparison between the asymptomatic vs moderate COVID-19 groups; AC vs HP, comparison between the asymptomatic and the hospitalized COVID-19 group; AP vs HP, comparison between the COVID-19 moderate and the hospitalized groups. (b) metrics of performance for each comparison group, left, predicted receiver operating characteristics (ROC) plots, showing the area under the curve (AUC) values, and right, predicted responses plots with the threshold value indicated as a red dash line.

**Table 2.**
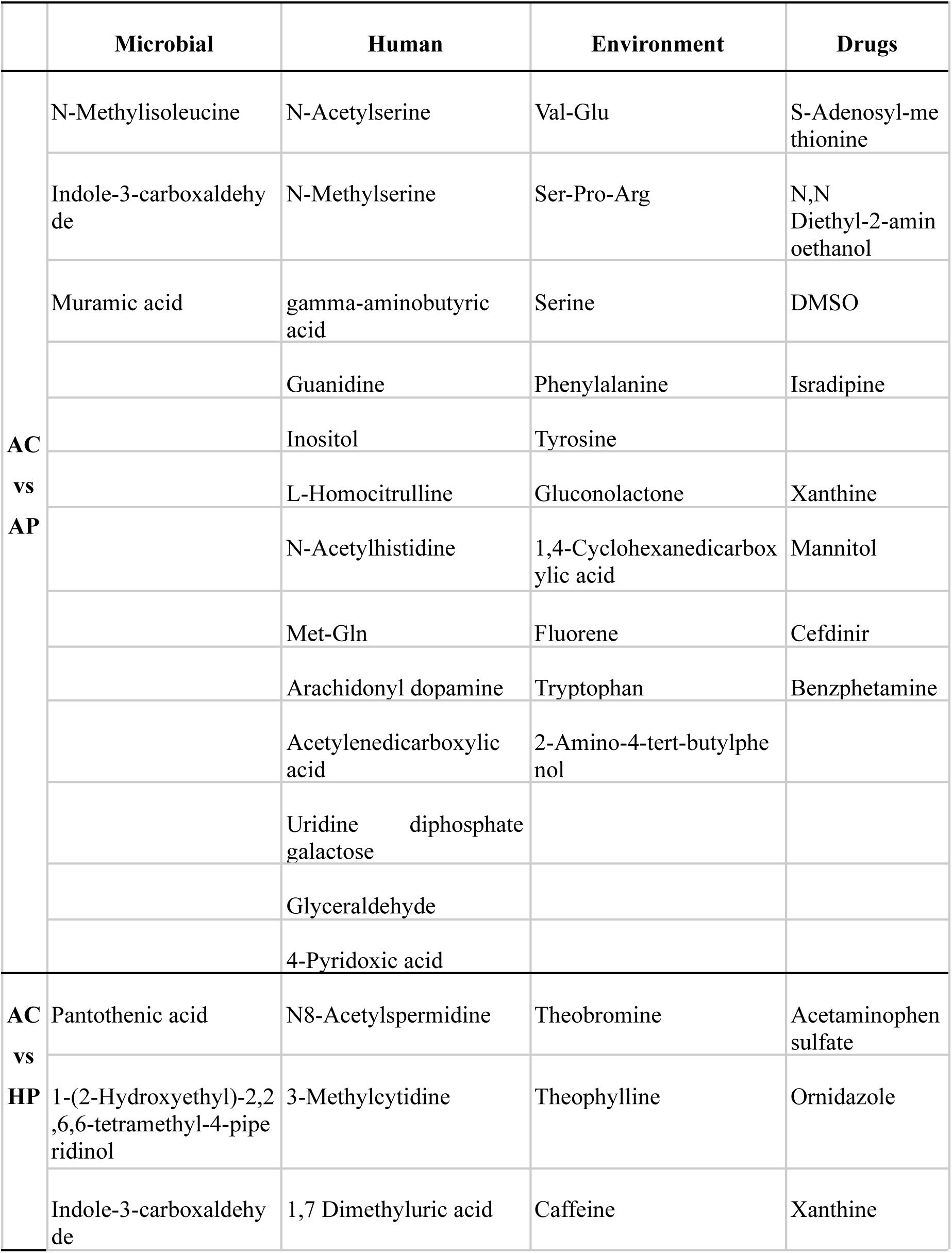
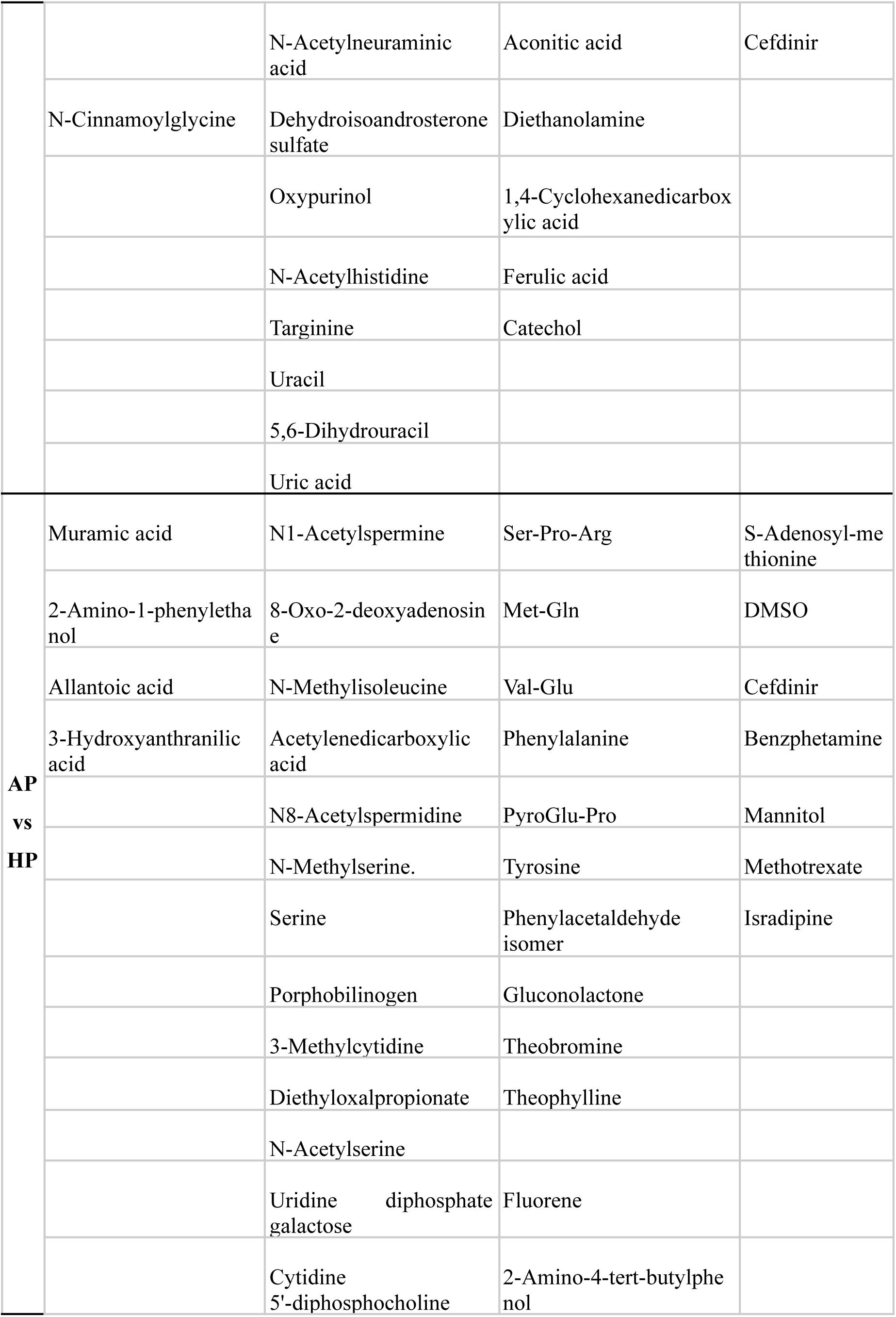
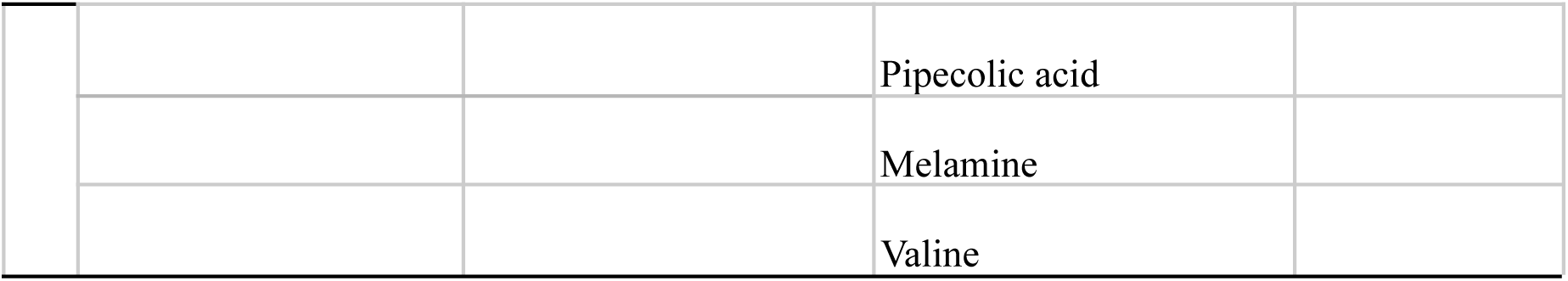
Origin of metabolites showing significant association with severity of COVID-19. Origin, microbial (M), environmental (E), human (H), drugs (D).

### Amino acid metabolites are strongly associated with moderate COVID cases and bacterial metabolites with severe cases

Infection with SARS-CoV-2 was associated with markedly significant changes of metabolites in saliva, particularly in moderate cases. The contribution of each metabolite to the identity of the groups was determined using a volcano plot (**Figure 3**). When comparing AP with the AC group, strongly significant increases in metabolites were observed. Notably, dipeptides, such as Val-Glu and Met-Gln showed highly significant increases in AP, with *p*-value exceeding 10^-40^ (Figure 3A). Additionally, aminoacids and their derivatives, including serine, phenylalanine, tyrosine, N-Acetylserine, N-Methyl Serine were also in a higher abundance in AP subjects with *p-*values higher than 10^-20^. In contrast, in the AC group Ser-Pro-Arg (*p*-value<10^-20^), S-Adenosyl-methionine and N-Methylisoleucine (*p*-values around 10^-10^) were significantly higher.

**Figure 3.**
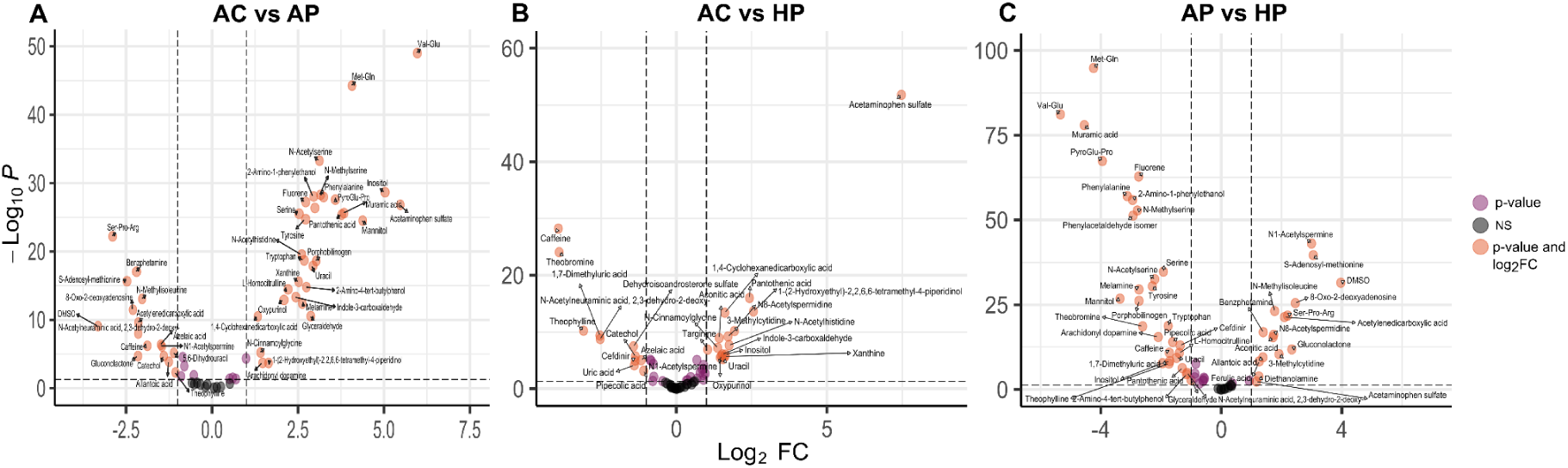
Pairwise differential abundance analysis of metabolites among the three clinical groups. Volcano plot shows metabolite fold changes (FC) on x-axis and the False Discovery Rate (FDR) on y-axis. Dashed lines indicate threshold, FC ≥ 1.0 (either positive or negative) and FDR > -log (0.05), metabolites in orange dots presented significant abundance. (A) AC vs AP comparison between the asymptomatic control group and the ambulatory SARS-CoV-2 positive group; (B) AC vs HP, comparison between the asymptomatic control group and the hospitalized SARS-CoV-2 positive group; (C) AP vs HP, comparison between ambulatory patients and the hospitalized SARS-CoV-2 positive group.

The metabolome of severe-hospitalized patients (HP) presented a rather different pattern when compared with AC (Figure 3B), and three bacteria-derived metabolites, pantothenic acid, 1-(2-Hydroxyethyl)-2,2,6,6-tetramethyl-4-piperidinol and indole-3-carboxaldehyde increased significantly. Also, human-derived N8-acetylspermidine and 3-methylcytidine concentrations were significantly higher, whereas amino acids or their derivatives were almost absent (as opposed to results in the AP group). In contrast, the saliva of AC individuals was significantly enriched with caffeine and its derivatives and with human-derived 1,7 dimethyluric acid, N-acetylneuraminic acid, and dehydroisoandrosterone sulfate.

We next explored the differences between the two SARS-CoV-2 infected patients, AP and HP (figure 3C), and observed that N1-acetylspermidine, 8-oxo-2-deoxyadenosine, S-adenosyl-methionine, and dimethyl-sulfoxide (DMSO) were significantly enriched in the HP group (10^-25^), it is a compound that is a solvent for drugs including antivirals. It is possible that its presence is due to the treatment regimen. Besides, N-methylisoleucine, Ser-Pro-Arg, acetylenedicarboxilic acid, and N8-acetylspermidine were also significantly more abundant in HP patients. In contrast, mostly amino acids and their derivatives were extremely higher in the AP group, with Met-Gln and Val-Glu havig p-values of 10^-80^, whereas phenylalanine, pyroGlu-Pro, and N-methylserine p-values of 10^-50^, and serine, N-acetylserine and tyrosine with p-values 10^-25^. Furthermore, muramic acid and 2-amino-1-phenylethanol, were two bacterial products highly enriched in AP (10^-50^).

Figure 4 illustrates the concentration of some metabolites selected because their concentration was significantly different between the clinical groups, in order to show the actual concentration differences between groups. Metabolites are presented according to their source, microbial, human or from the environment.

**Figure 4.**
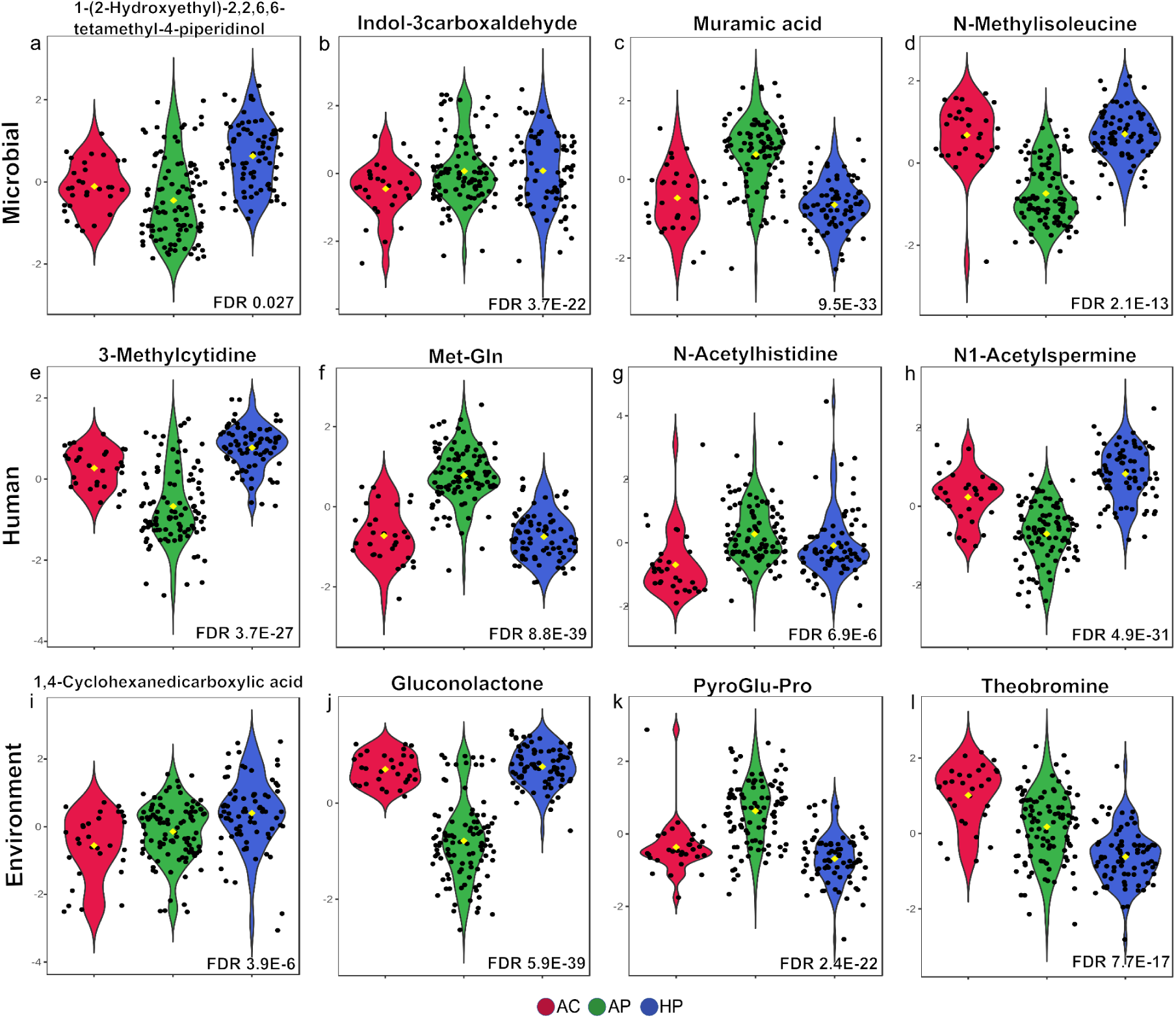
Violin plots of metabolites with differences among the AC, AP and HP groups, according to their source. A selection of metabolites for three categories (microbial, human, environment) is shown, False discovery Rate (FDR) values (< 0.05) from a one-way ANOVA test are indicated.

### Some metabolites distinguish deceased from severe-hospitalized patients

In the group of hospitalized patients, there were 17 deceased patients (DHP) and we decided to search for any difference between these 17 and the other hospitalized patients (HP). S-Adenosyl-methionine, N-acetylhistidine, porphobilinogen, tyrosine, and bacteria-derived muramic acid were significantly increased in DHP compared to those in the HP group (**Supplemental** figure 2). Caffeine and its metabolites, theobromine and theophylline were notably more prevalent in DHP. In contrast, only 2-amino-4 tert.butylphenol was significantly increased in the HP group.

## Discussion

While several metabolome studies in COVID-19 patients have been published, most reported on a limited number of metabolites in plasma (Ansone et al. 2021) (Bourgin et al. 2021). The few studies conducted in saliva included a small number of patients with a more restricted search of metabolites (Frampas et al. 2021) (Pozzi et al. 2022) (Costa Dos Santos Junior et al. 2020). The present work is a comprehensive analysis of metabolites present in the saliva of patients with COVID-19 that shows significant changes in the composition and concentration of metabolites in patients infected with SARS-CoV-2 and specific metabolite patterns associated with the severity of disease. We will discuss our work in the context of both, the utility of our findings in the search of biomarkers but also in the search of knowledge to better understand pathogenic mechanisms and probable targets of this worrisome infection.

The presence of dipeptides Val-Glu and Met-Gln in saliva of patients with moderate COVID disease was outstandingly higher than in asymptomatic uninfected individuals. We could not find reports of these peptides in saliva, but saliva is known to contain several proteolytic enzymes (Thomadaki et al. 2011) and peptides might be the products of the activity of proteases. The marked abundance of only these two dipeptides would suggest the activity of specific proteases on specific proteins, and this might suggest Val-Glu and Met-Gln are a useful marker for SARS-CoV-2 infection in mild-moderate COVID episodes. Previous studies suggest saliva contains a mixture of enzymes resilient to protease inhibitors but with a tightly controlled activity that has been associated with health and disease (Garreto et al. 2021) (Turk 2006). Some dipeptides may present cell-signaling effects (https://hmdb.ca/metabolites/HMDB0029125) or regulatory functions important for health (Minen, Thirumalaikumar, and Skirycz 2023).

Phenylalanine, serine, tyrosine, and tryptophan were also increased in cases of mild-moderate disease, and may result from proteolytic activity, but may also be of microbial origin. Recent studies have shown that tryptophan catabolites produced by microbial members may have relevant roles in health and disease, they may stimulate the immune system, epithelial barrier, and even gut hormone secretion (Roager and Licht 2018). In contrast to our results, an exploratory study by Frampas et al. reports a decrease of phenylalanine and tyrosine in the saliva of infected COVID patients (Frampas et al. 2022). Patients in our study were recruited during the first wave of the pandemic, before vaccination, and this might be an important difference from other reports. In any case, it is important to explore which proteases are activated during SARS-CoV-2 infection and the role their products may have in the pathogenesis of COVID-19.

Two acetylated amino acids were strongly associated with mild-moderate COVID-19, N-acetylserine and N-acetylhistidine. These two acetylated amino acids have been reported to be elevated in sera of patients with chronic kidney disease and have been suggested as markers of tubular renal function (Wen et al. 2022) (Luo et al. 2020). In addition, N-acetylhistidine was found significantly increased during ischemia in pigs with induced myocardial infarction (Goetzman et al. 2022) and has been reported significantly decreased in the brains of Alzheimer’s disease patients (Hammond et al. 2021). N-terminal acetylation of amino acids has been suggested as part of the response to stress and is associated with cancer and other developmental disorders (Ree, Varland, and Arnesen 2018). In previous studies, acetylated amino acids have been detected mostly in sera. Our study shows for the first time that they can also be detected in saliva during SARS-CoV-2 infection, a virus that has shown an extraordinary ability to infect cells of several organs, including kidney and hearth, posing patients under a strong stress. Saliva N-acetylserine and N-acetylhistidine seem to represent sensitive markers of stress and probably a marker for patients at risk of developing more severe COVID-19 disease.

When compared with asymptomatic adults, muramic acid, a component of the cell wall in bacteria, was significantly increased in mild-moderate cases, suggesting that there might be an overgrowth of bacteria as a response to SARS-CoV-2 infection. Indole-3-carboxaldehyde, a bacterial metabolite of tryptophan, also increased in mild-moderate cases. This metabolite is synthesized particularly by *Lactobacillus spp*. and is known to stimulate IL-22 production and increase the immune reactivity of mucosas (Zelante et al. 2013). The increase in these two metabolites attest to relevant changes in the oral microbiota of patients infected with SARS-CoV-2 infection.

In this work Ser-Pro-Arg was significantly reduced in mild-moderate patients, which is probably a product of proteolytic activity. We did not find reports of this peptide in saliva or other human fluids, although this tripeptide sequence is present in at least five repeats in the histone H1 and they might be important for the structure of the protein (Strickland et al. 1980). S-Adenosyl-methionine (SAM), a key metabolite in methylation reactions, was significantly reduced in mild-moderate patients, which suggests that it is consumed during infection. SAM is produced in the liver and has been associated with liver disease, including steatosis, hepatocarcinoma, or hepatomegaly and might be a sensitive marker for liver damage, one of the multiple targets of SARS-CoV-2 (Pascale et al. 2022). It is interesting to note that the administration of SAM to patients with HCV inhibits the expression of the virus and improves the activity of peginterferon (Pascale et al. 2022). N-methylisoleucine was also reduced in mild-moderate patients. This metabolite has been found upregulated in the plasma of patients with glioblastoma (Aboud et al. 2023), although its role during viral infections is poorly understood. It is tempting to suggest that Ser-Pro-Arg, SAM and N-methylisoleucine are amino acid metabolites that are significantly consumed (reduced) during acute SARS-CoV-2 infection to prevent severe disease.

We next examined differences between asymptomatic and severe-hospitalized patients. As a result, we identified two bacterial metabolites among the most significantly increased in the hospitalized patients: pantothenic acid and Indole-3-carboxaldehyde (Figure 3b). Pantothenic acid is a precursor in the synthesis of CoA and genomic studies have revealed that many Bacteroidetes and Proteobacteria produce CoA (Magnúsdóttir et al. 2015). Indole is produced by many bacteria and multiple physiological functions have been described, including plasmid stability, drug resistance control or biofilm formation (Lee and Lee 2010). Probably the observed increase of these two metabolites in severe cases reflects a response of bacteria to changes in their microenvironment.

Polyamines are essential for cell survival, participate in multiple processes like metabolism or regulation of replication, and are undoubtedly associated with multiple diseases (Wallace, Fraser, and Hughes 2003). N8-acetylspermidine is a polyamine produced in episodes of cardiac ischemia, cancer, or even infections and is usually detected in plasma (Nayak et al. 2020). A study in patients with gastric cancer reported that increased plasma levels of acetylated polyamines correlated with severe cases that required hospitalization (Bourgin et al. 2021). Here we found increased levels in the saliva of patients with severe COVID-19 infection, most probably a marker of alert for the extensive damage to multiple organs.

Whereas the function of 1-(2-Hydroxyethyl)-2,2,6,6-tetramethyl-4-piperidinol remains unclear, it is not a human metabolite and is found only in the blood of people exposed to a possible source such as additive stabilizer in the synthesis of plastics and of light stabilizer additives. This compound was increased in serum of rats exposed to stress, and authors suggested it may have a role in the gut-brain axis dialogue (Qu et al. 2023). In addition, 1-4-cyclohexanedicarboxilic acid is a polyvinyl chloride used as a flavor in food and beverage and with medical applications (Wenzel et al. 2021) that was also increased in severe cases. The increased level of these 2 metabolites in the saliva of hospitalized patients may suggest that previous exposure to these compounds represents a risk of developing severe COVID-19 disease.

3-methylcytidine also increased in severe-hospitalized patients. It is a key regulator of transcription, responsible for fine-tuning gene expression in response to changing conditions, and considered an epitranscriptome marker (Bohnsack et al. 2022). It is possible that in severe COVID-19 there is a demand for 3-methylcytidine because of a need for increased transcription activity, particularly requiring tRNA m3-C32 modifications.

Learning the differences in metabolites between moderate and severe cases was also important. Interestingly, N1-acetylspermine was the most significantly enriched metabolite in severe cases, an intermediate in the conversion of spermine to spermidine. Polyamines are important in key cellular processes like growth and differentiation and have been reported to be associated with cardiac ischemia and pancreatic or breast cancer (Asai et al. 2018) and could be useful as markers for severe organ damage, as observed in severe COVID-19 cases.

It was intriguing to note the striking differences in metabolites between moderate and severe cases. For instance, several metabolites that significantly decreased in moderate patients when compared with asymptomatic (including 8-oxo-2-deoxyadenosine, S-adenosyl-methionine, N-methylisoleucine, Ser-Pro-Arg, acetylenedicarboxylic acid and gluconolactone) they showed values similar or higher to the asymptomatic individuals in the severe cases. Furthermore, the strong dipeptide and amino acid response observed in mild-moderate cases was absent in the severe cases. It is tempting to speculate that the stronger metabolic changes observed in moderate cases represent an intense response that prevents patients from developing severe disease. In contrast, in severe patients the strong increase was in alarm metabolites like N-acetylspermine, S-adenosyl--methionine and 8-oxo-2-deoxyadenosine, already described as markers of other severe diseases.

Finally, it is interesting to note that when compared the deceased cases with the severe patients the most significant difference was a high increase of N-acetylhistidine in fatal cases, a metabolite that has been associated with kidney failure (Luo et al. 2020) but also myocardial ischemia (Goetzman et al. 2022) and even Alzheimer disease (Hammond et al. 2021). Porphobilinogen was also increased in deceased cases, and urinary porphobilinogen has been suggested as a marker for hepatic porphyrias but it is also relevant in kidney disease (Aarsand, Petersen, and Sandberg 2006). Thus, the detection of N-acetylhistidine and porphobilinogen in saliva may be markers of kidney, heart or brain damage in severe COVID-19 cases with increased risk of a fatal outcome.

In conclusion, this study documents strongly significant changes in the metabolome of saliva of patients infected with SARS-CoV-2 presenting a moderate COVID-19 disease, particularly in dipeptides and amino acids, probably because of specific proteases. Metabolome changes in severe cases were also highly significant, but with a pattern completely different, where metabolites previously associated with damage to kidney, hearth, brain and liver were significantly increased. Some of these markers were further increased in deceased patients. This study shows the convenience of saliva for exhaustive analyses of metabolic profiles in patients with COVID-19 and suggests its potential utility for diagnosis, prognosis and even pathogenesis studies.

## References

Aarsand, Aasne K., Per Hyltoft Petersen, and Sverre Sandberg. 2006. “Estimation and Application of Biological Variation of Urinary Delta-Aminolevulinic Acid and Porphobilinogen in Healthy Individuals and in Patients with Acute Intermittent Porphyria.” Clinical Chemistry 52 (4): 650–56.

Aboud, Orwa, Yin Liu, Lina Dahabiyeh, Ahmad Abuaisheh, Fangzhou Li, John Paul Aboubechara, Jonathan Riess, et al. 2023. “Profile Characterization of Biogenic Amines in Glioblastoma Patients Undergoing Standard-of-Care Treatment.” Biomedicines 11 (8). 10.3390/biomedicines11082261.

Ansone, Laura, Monta Briviba, Ivars Silamikelis, Anna Terentjeva, Ingus Perkons, Liga Birzniece, Vita Rovite, et al. 2021. “Amino Acid Metabolism Is Significantly Altered at the Time of Admission in Hospital for Severe COVID-19 Patients: Findings from Longitudinal Targeted Metabolomics Analysis.” Microbiology Spectrum 9 (3): e0033821.

Asai, Yasutsugu, Takao Itoi, Masahiro Sugimoto, Atsushi Sofuni, Takayoshi Tsuchiya, Reina Tanaka, Ryosuke Tonozuka, et al. 2018. “Elevated Polyamines in Saliva of Pancreatic Cancer.” Cancers 10 (2). 10.3390/cancers10020043.

Blaženović, Ivana, Tobias Kind, Michael R. Sa, Jian Ji, Arpana Vaniya, Benjamin Wancewicz, Bryan S. Roberts, et al. 2019. “Structure Annotation of All Mass Spectra in Untargeted Metabolomics.” Analytical Chemistry 91 (3): 2155–62.

Bohnsack, Katherine E., Nicole Kleiber, Nicolas Lemus-Diaz, and Markus T. Bohnsack. 2022. “Roles and Dynamics of 3-Methylcytidine in Cellular RNAs.” Trends in Biochemical Sciences 47 (7): 596–608.

Bourgin, Mélanie, Lisa Derosa, Carolina Alves Costa Silva, Anne-Gaëlle Goubet, Agathe Dubuisson, François-Xavier Danlos, Claudia Grajeda-Iglesias, et al. 2021. “Circulating Acetylated Polyamines Correlate with Covid-19 Severity in Cancer Patients.” Aging 13 (17): 20860–85.

Costa Dos Santos Junior, Gilson, Claudia Maria Pereira, Tatiana Kelly da Silva Fidalgo, and Ana Paula Valente. 2020. “Saliva NMR-Based Metabolomics in the War Against COVID-19.” Analytical Chemistry 92 (24): 15688–92.

Dewhirst, Floyd E., Tuste Chen, Jacques Izard, Bruce J. Paster, Anne C. R. Tanner, Wen-Han Yu, Abirami Lakshmanan, and William G. Wade. 2010. “The Human Oral Microbiome.” Journal of Bacteriology 192 (19): 5002–17.

Fan, Sili, Tobias Kind, Tomas Cajka, Stanley L. Hazen, W. H. Wilson Tang, Rima Kaddurah-Daouk, Marguerite R. Irvin, Donna K. Arnett, Dinesh K. Barupal, and Oliver Fiehn. 2019. “Systematic Error Removal Using Random Forest for Normalizing Large-Scale Untargeted Lipidomics Data.” Analytical Chemistry 91 (5): 3590–96.

Frampas, Cecile F., Katie Longman, Matt Spick, Holly-May Lewis, Catia D. S. Costa, Alex Stewart, Deborah Dunn-Walters, et al. 2022. “Untargeted Saliva Metabolomics by Liquid Chromatography-Mass Spectrometry Reveals Markers of COVID-19 Severity.” PloS One 17 (9): e0274967.

Frampas, Cecile F., Katie Longman, Matt P. Spick, Holly M. Lewis, Catia D. S. Costa, Alex Stewart, Deborah Dunn-Walters, et al. 2021. “Untargeted Saliva Metabolomics Reveals COVID-19 Severity.” bioRxiv. medRxiv. 10.1101/2021.07.06.21260080.

Garreto, Laís, Sébastien Charneau, Samuel Coelho Mandacaru, Otávio T. Nóbrega, Flávia N. Motta, Carla N. de Araújo, Audrey C. Tonet, et al. 2021. “Mapping Salivary Proteases in Sjögren’s Syndrome Patients Reveals Overexpression of Dipeptidyl Peptidase-4/CD26.” Frontiers in Immunology 12 (June):686480.

Goetzman, Eric, Zhenwei Gong, Dhivyaa Rajasundaram, Ishan Muzumdar, Traci Goodchild, David Lefer, and Radhika Muzumdar. 2022. “Serum Metabolomics Reveals Distinct Profiles during Ischemia and Reperfusion in a Porcine Model of Myocardial Ischemia-Reperfusion.” International Journal of Molecular Sciences 23 (12). 10.3390/ijms23126711.

Hammond, Tyler C., Xin Xing, Lucy M. Yanckello, Arnold Stromberg, Ya-Hsuan Chang, Peter T. Nelson, and Ai-Ling Lin. 2021. “Human Gray and White Matter Metabolomics to Differentiate APOE and Stage Dependent Changes in Alzheimer’s Disease.” Journal of Cellular Immunology 3 (6): 397–412.

Harari, Sheri, Danielle Miller, Shay Fleishon, David Burstein, and Adi Stern. 2024. “Using Big Sequencing Data to Identify Chronic SARS-Coronavirus-2 Infections.” Nature Communications 15 (1): 648.

Larios Serrato, Violeta, Beatriz Meza, Carolina Gonzalez-Torres, Javier Gaytan-Cervantes, Joaquín González Ibarra, Clara Esperanza Santacruz Tinoco, Yu-Mei Anguiano Hernández, et al. 2023. “Diversity, Composition, and Networking of Saliva Microbiota Distinguish the Severity of COVID-19 Episodes as Revealed by an Analysis of 16S rRNA Variable V1-V3 Region Sequences.” mSystems 8 (4): e0106222.

Lee, Jin-Hyung, and Jintae Lee. 2010. “Indole as an Intercellular Signal in Microbial Communities.” FEMS Microbiology Reviews 34 (4): 426–44.

Love, Michael I., Wolfgang Huber, and Simon Anders. 2014. “Moderated Estimation of Fold Change and Dispersion for RNA-Seq Data with DESeq2.” Genome Biology 15 (12): 550.

Luo, Shengyuan, Aditya Surapaneni, Zihe Zheng, Eugene P. Rhee, Josef Coresh, Adriana M. Hung, Girish N. Nadkarni, et al. 2020. “Variants, N-Acetylated Amino Acids, and Progression of CKD.” Clinical Journal of the American Society of Nephrology: CJASN 16 (1): 37–47.

Magnúsdóttir, Stefanía, Dmitry Ravcheev, Valérie de Crécy-Lagard, and Ines Thiele. 2015. “Systematic Genome Assessment of B-Vitamin Biosynthesis Suggests Co-Operation among Gut Microbes.” Frontiers in Genetics 6 (April):148.

Minen, Romina Ines, Venkatesh P. Thirumalaikumar, and Aleksandra Skirycz. 2023. “Proteinogenic Dipeptides, an Emerging Class of Small-Molecule Regulators.” Current Opinion in Plant Biology 75 (102395): 102395.

Mollentze, Nardus, and Daniel G. Streicker. 2020. “Viral Zoonotic Risk Is Homogenous among Taxonomic Orders of Mammalian and Avian Reservoir Hosts.” Proceedings of the National Academy of Sciences of the United States of America 117 (17): 9423–30.

Moreno, Elena, Sergio Ciordia, Santos Milhano Fátima, Daniel Jiménez, Javier Martínez-Sanz, Pilar Vizcarra, Raquel Ron, et al. 2024. “Proteomic Snapshot of Saliva Samples Predicts New Pathways Implicated in SARS-CoV-2 Pathogenesis.” Clinical Proteomics 21 (1): 37.

Nayak, Aditi, Chang Liu, Anurag Mehta, Yi-An Ko, Ayman S. Tahhan, Devinder S. Dhindsa, Karan Uppal, et al. 2020. “N8-Acetylspermidine: A Polyamine Biomarker in Ischemic Cardiomyopathy With Reduced Ejection Fraction.” Journal of the American Heart Association 9 (11): e016055.

Pascale, Rosa M., Maria M. Simile, Diego F. Calvisi, Claudio F. Feo, and Francesco Feo. 2022. “S-Adenosylmethionine: From the Discovery of Its Inhibition of Tumorigenesis to Its Use as a Therapeutic Agent.” Cells 11 (3). 10.3390/cells11030409.

Pozzi, Chiara, Riccardo Levi, Daniele Braga, Francesco Carli, Abbass Darwich, Ilaria Spadoni, Bianca Oresta, et al. 2022. “A ‘Multiomic’ Approach of Saliva Metabolomics, Microbiota, and Serum Biomarkers to Assess the Need of Hospitalization in Coronavirus Disease 2019.” Gastro Hep Advances 1 (2): 194–209.

Qu, Youge, Akifumi Eguchi, Li Ma, Xiayun Wan, Chisato Mori, and Kenji Hashimoto. 2023. “Role of the Gut-Brain Axis via the Subdiaphragmatic Vagus Nerve in Stress Resilience of 3,4-Methylenedioxymethamphetamine in Mice Exposed to Chronic Restrain Stress.” Neurobiology of Disease 189 (December):106348.

Ree, Rasmus, Sylvia Varland, and Thomas Arnesen. 2018. “Spotlight on Protein N-Terminal Acetylation.” Experimental & Molecular Medicine 50 (7): 1–13.

Roager, Henrik M., and Tine R. Licht. 2018. “Microbial Tryptophan Catabolites in Health and Disease.” Nature Communications 9 (1): 3294.

Sievers, Benjamin L., Mark T. K. Cheng, Kata Csiba, Bo Meng, and Ravindra K. Gupta. 2024. “SARS-CoV-2 and Innate Immunity: The Good, the Bad, and the ‘Goldilocks.’” Cellular & Molecular Immunology 21 (2): 171–83.

Strickland, W. N., M. Strickland, P. C. de Groot, C. Von Holt, and B. Wittmann-Liebold. 1980. “The Primary Structure of Histone H1 from Sperm of the Sea Urchin Parechinus Angulosus. 1. Chemical and Enzymatic Fragmentation of the Protein and the Sequence of Amino Acids in the Four N-Terminal Cyanogen Bromide Peptides.” European Journal of Biochemistry / FEBS 104 (2): 559–66.

Tang, Yujun, Jiajia Liu, Dingyi Zhang, Zhenghao Xu, Jinjun Ji, and Chengping Wen. 2020. “Cytokine Storm in COVID-19: The Current Evidence and Treatment Strategies.” Frontiers in Immunology 11 (July):1708.

Thomadaki, K., E. J. Helmerhorst, N. Tian, X. Sun, W. L. Siqueira, D. R. Walt, and F. G. Oppenheim. 2011. “Whole-Saliva Proteolysis and Its Impact on Salivary Diagnostics.” Journal of Dental Research 90 (11): 1325–30.

Tsang, Tim K., Kyu Han Lee, Betsy Foxman, Angel Balmaseda, Lionel Gresh, Nery Sanchez, Sergio Ojeda, et al. 2020. “Association Between the Respiratory Microbiome and Susceptibility to Influenza Virus Infection.” Clinical Infectious Diseases: An Official Publication of the Infectious Diseases Society of America 71 (5): 1195–1203.

Tsugawa, Hiroshi, Tomas Cajka, Tobias Kind, Yan Ma, Brendan Higgins, Kazutaka Ikeda, Mitsuhiro Kanazawa, Jean VanderGheynst, Oliver Fiehn, and Masanori Arita. 2015. “MS-DIAL: Data-Independent MS/MS Deconvolution for Comprehensive Metabolome Analysis.” Nature Methods 12 (6): 523–26.

Turk, Boris. 2006. “Targeting Proteases: Successes, Failures and Future Prospects.” Nature Reviews. Drug Discovery 5 (9): 785–99.

Wallace, Heather M., Alison V. Fraser, and Alun Hughes. 2003. “A Perspective of Polyamine Metabolism.” Biochemical Journal 376 (Pt 1): 1–14.

Wen, Donghai, Zihe Zheng, Aditya Surapaneni, Bing Yu, Linda Zhou, Wen Zhou, Dawei Xie, et al. 2022. “Metabolite Profiling of CKD Progression in the Chronic Renal Insufficiency Cohort Study.” JCI Insight 7 (20). 10.1172/jci.insight.161696.

Wenzel, Abby G., Jessica L. Reiner, Satomi Kohno, Bethany J. Wolf, John W. Brock, Lori Cruze, Roger B. Newman, and John R. Kucklick. 2021. “Biomonitoring of Emerging DINCH Metabolites in Pregnant Women in Charleston, SC: 2011-2014.” Chemosphere 262 (January):128369.

Wickham, Hadley. 2016. Ggplot2: Elegant Graphics for Data Analysis. PDF. 2nd ed. Use R! Cham, Switzerland: Springer International Publishing.

Zelante, Teresa, Rossana G. Iannitti, Cristina Cunha, Antonella De Luca, Gloria Giovannini, Giuseppe Pieraccini, Riccardo Zecchi, et al. 2013. “Tryptophan Catabolites from Microbiota Engage Aryl Hydrocarbon Receptor and Balance Mucosal Reactivity via Interleukin-22.” Immunity 39 (2): 372–85.

