## Supplemental Table 1 for "Composition of saliva metabolome is significantly associated with SARS-CoV2 infection and with severity of COVID-19 disease"

**Supplemental Table 2. Differences in intensity of the selected metabolites by group.**
Origin: microbial (M), environmental (E), human (H), drugs (D).

| Annotation | Species | MSI Level | m/z | ESI mode | RT | FDR | VIP | Origin |
|---|---|---|---|---|---|---|---|---|
| 1-(2-Hydroxyethyl)-2,2,6,6-tetramethyl-4-piperidinol | [M+H]+ | 2 | 202.17 | ESI pos | 1.02 | 2.70E-02 | AC vs HP | M |
| 1,4-Cyclohexanedicarboxylic acid | [M-H]- | 3 | 171.06 | ESI neg | 0.49 | 3.90E-06 | AC vs HP | E |
| 1,7-Dimethyluric acid | [M-H]- | 3 | 195.05 | ESI neg | 0.89 | 1.73E-08 | AC vs HP | H |
| 2-Amino-1-phenylethanol | [M+H-H2O]+ | 2 | 120.08 | ESI pos | 1.70 | 1.13E-24 | AP vs HP | M |
| 2-Amino-4-tert-butylphenol | [M+H-H2O]+ | 2 | 148.11 | ESI pos | 0.56 | 9.26E-05 | AC vs AP, AC vs HP | E |
| 3-Hydroxyanthranilic acid | [M-H]- | 3 | 152.03 | ESI neg | 0.23 | 3.73E-11 | AC vs HP | M |
| 3-Methylcytidine | [M+H]+ | 3 | 258.10 | ESI pos | 1.75 | 3.70E-27 | AC vs HP, AC vs HP | H |
| 4-Pyridoxic acid | [M-H]- | 3 | 182.04 | ESI neg | 0.49 | 2.90E-03 | AC vs AP | H |
| 5,6-Dihydrouracil | [M+H]+ | 3 | 115.05 | ESI pos | 0.35 | 1.02E-10 | AC vs HP | H |
| 8-Oxo-2-deoxyadenosine | [M+H]+ | 3 | 268.10 | ESI pos | 1.14 | 3.46E-21 | AC vs HP | H |
| Acetaminophen sulfate | [M-H]- | 2 | 230.01 | ESI neg | 0.35 | 1.17E-08 | AC vs HP | D |
| Acetylenedicarboxylic acid | [M-H]- | 3 | 112.98 | ESI neg | 1.90 | 2.88E-45 | AC vs AP, AC vs HP | H |
| Aconitic acid | [M-H]- | 3 | 173.00 | ESI neg | 1.97 | 1.20E-20 | AC vs HP | E |
| Allantoic acid | [M-H]- | 3 | 175.04 | ESI neg | 2.11 | 1.58E-27 | AC vs HP | M |
| Arachidonyl dopamine | [M-H]- | 3 | 438.29 | ESI neg | 0.34 | 0.7639 | AC vs AP | H |
| Azelaic acid | [M-H]- | 2 | 187.09 | ESI neg | 0.34 | 8.16E-12 | AC vs HP | E |
| Benzphetamine | [M+H]+ | 3 | 240.17 | ESI pos | 0.24 | 5.99E-33 | AC vs AP, AC vs HP | D |

| | | | | | | | | |
|---|---|---|---|---|---|---|---|---|
| Caffeine | [M+H]+ | 1 | 195.08 | ESI pos | 0.26 | 1.91E-10 | AC vs HP | E |
| Catechol | [M-H]- | 3 | 109.03 | ESI neg | 0.26 | 6.33E-12 | AC vs HP | E |
| Cefdinir | [M-H]- | 3 | 394.02 | ESI neg | 1.74 | 1.79E-17 | AC vs AP, AC vs HP, AC vs HP | D |
| Cytidine 5'-diphosphocholine | [M+H]+ | 2 | 489.10 | ESI pos | 2.48 | 1.72E-13 | AC vs HP | H |
| Dehydroisoandrosterone sulfate | [M-H]- | 3 | 367.15 | ESI neg | 0.33 | 5.30E-11 | AC vs HP | H |
| Diethanolamine | [M+H]+ | 2 | 106.08 | ESI pos | 1.74 | 6.80E-16 | AC vs HP | E |
| Diethyloxalpropionate | [M-H]- | 3 | 201.07 | ESI neg | 0.74 | 4.03E-08 | AC vs HP | H |
| DMSO | [M+H]+ | 1 | 79.02 | ESI pos | 0.33 | 7.02E-40 | AC vs AP, AC vs HP | D |
| Ferulic acid | [M-H]- | 3 | 193.05 | ESI neg | 0.35 | 1.00E-18 | AC vs HP | E |
| Fluorene | [M+H]+ | 3 | 167.08 | ESI pos | 1.70 | 4.87E-26 | AC vs AP, AC vs HP | E |
| gamma-aminobutyric acid | [M+H]+ | 3 | 104.07 | ESI pos | 1.92 | 0.1233 | AC vs AP | H |
| Gluconolactone | [M+H]+ | 3 | 179.05 | ESI pos | 1.96 | 5.90E-39 | AC vs AP, AC vs HP | E |
| Glyceraldehyde | [M-H]- | 3 | 89.02 | ESI neg | 1.92 | 0.5423 | AC vs AP | H |
| Guanidine | [M+H]+ | 2 | 60.05 | ESI pos | 1.52 | 1.72E-22 | AC vs AP, AC vs HP | H |
| Indole-3-carboxaldehyde | [M-H]- | 3 | 144.04 | ESI neg | 0.34 | 3.70E-22 | AC vs AP, AC vs HP | M |
| Inositol | [M-H]- | 3 | 179.05 | ESI neg | 1.92 | 4.49E-06 | AC vs AP | H |
| Isradipine | [M-H]- | 3 | 370.14 | ESI neg | 0.23 | 1.02E-04 | AC vs AP, AC vs HP | D |
| L-Homocitrulline | [M+H]+ | 2 | 190.11 | ESI pos | 2.13 | 1.36E-07 | AC vs AP | H |
| Mannitol | [M-H]- | 3 | 181.07 | ESI neg | 1.92 | 1.67E-14 | AC vs AP, AC vs HP | D |

| | | | | | | | | |
|---|---|---|---|---|---|---|---|---|
| Melamine | [M+H]+ | 2 | 127.07 | ESI pos | 1.08 | 1.09E-04 | AC vs HP | E |
| Met-Gln | [M+H]+ | 2 | 278.12 | ESI pos | 2.03 | 8.80E-39 | AC vs AP, AC vs HP | H |
| Methotrexate | [M-H]- | 3 | 453.16 | ESI neg | 1.93 | 7.16E-18 | AC vs HP | D |
| Muramic acid | [M-H]- | 3 | 250.09 | ESI neg | 2.02 | 9.50E-33 | AC vs AP, AC vs HP | M |
| N,N-Diethyl-2-aminoethanol | [M+H]+ | 2 | 118.12 | ESI pos | 0.85 | 8.34E-33 | AC vs AP | D |
| N1-Acetylspermine | [M+H]+ | 2 | 245.23 | ESI pos | 2.48 | 4.90E-31 | AC vs HP | H |
| N8-Acetylspermidine | [M+H]+ | 3 | 188.17 | ESI pos | 2.17 | 1.80E-12 | AC vs HP | H |
| N-Acetylhistidine | [M+H]+ | 1 | 198.08 | ESI pos | 1.98 | 6.90E-06 | AC vs AP, AC vs HP | H |
| N-Acetylneuraminic acid, 2,3-dehydro-2-deoxy- | [M+Na]+ | 2 | 314.08 | ESI pos | 2.07 | 1.06E-09 | AC vs HP | H |
| N-Acetylserine | [M-H]- | 3 | 146.04 | ESI neg | 1.92 | 1.20E-20 | AC vs AP, AC vs HP | H |
| N-Cinnamoylglycine | [M-H]- | 3 | 204.06 | ESI neg | 0.71 | 5.01E-04 | AC vs HP | M |
| N-Methylisoleucine | [M+H]+ | 3 | 146.11 | ESI pos | 1.63 | 2.10E-13 | AC vs AP, AC vs HP | M, H |
| N-Methylserine | [M+H]+ | 1 | 120.06 | ESI pos | 2.01 | 2.44E-24 | AC vs AP, AC vs HP | H |
| Ornidazole | [M+H]+ | 3 | 220.05 | ESI pos | 0.25 | 6.57E-04 | AC vs HP | D |
| Oxypurinol | [M-H]- | 2 | 151.02 | ESI neg | 1.39 | 4.03E-07 | AC vs HP | H |
| Pantothenic acid | [M+H]+ | 2 | 220.11 | ESI pos | 1.20 | 1.44E-06 | AC vs HP | M |
| Phenylacetaldehyde isomer | [M+H-H2 O]+ | 2 | 103.05 | ESI pos | 1.70 | 1.11E-22 | AC vs HP | E |
| Phenylalanine | [M-H]- | 3 | 164.07 | ESI neg | 1.70 | 2.26E-23 | AC vs HP | E |
| Pipecolic acid | [M+H]+ | 3 | 130.08 | ESI pos | 1.92 | 1.40E-10 | AC vs HP | M, H |
| Porphobilinogen | [M-H]- | 1 | 225.08 | ESI neg | 1.89 | 2.23E-20 | AC vs HP | H |

| | | | | | | | | |
|---|---|---|---|---|---|---|---|---|
| PyroGlu-Pro | [M+H]+ | 2 | 227.10 | ESI pos | 1.89 | 2.40E-22 | AC vs HP | E |
| S-Adenosyl-methionine | [M+H]+ | 1 | 399.13 | ESI pos | 2.44 | 5.58E-21 | AC vs AP, AC vs HP | D |
| Serine | [M-H]- | 3 | 104.03 | ESI neg | 2.11 | 4.18E-18 | AC vs AP | H, E |
| Ser-Pro-Arg | [M+H]+ | 2 | 359.20 | ESI pos | 2.42 | 4.07E-14 | AC vs AP | E |
| Targinine | [M+H]+ | 1 | 189.13 | ESI pos | 2.26 | 5.31E-03 | AC vs HP | H |
| Theobromine | [M+H]+ | 2 | 181.07 | ESI pos | 0.34 | 7.70E-17 | AC vs HP, AC vs HP | E |
| Theophylline | [M-H]- | 3 | 179.05 | ESI neg | 0.33 | 4.02E-16 | AC vs HP, AC vs HP | E |
| Tryptophan | [M-H]- | 3 | 203.08 | ESI neg | 1.68 | 1.67E-14 | AC vs AP, AC vs HP | E |
| Tyrosine | [M+H]+ | 1 | 182.07 | ESI pos | 1.91 | 6.30E-18 | AC vs HP | E |
| Uracil | [M+H]+ | 1 | 113.03 | ESI pos | 0.50 | 1.09E-09 | AC vs HP | H |
| Uric acid | [M-H]- | 2 | 167.02 | ESI neg | 1.92 | 9.58E-06 | AC vs HP | H |
| Uridine diphosphate galactose | [M-H]- | 3 | 565.05 | ESI neg | 2.53 | 5.66E-22 | AC vs AP, AC vs HP | H |
| Val-Glu | [M-H]- | 2 | 245.11 | ESI neg | 2.03 | 6.22E-22 | AC vs HP | E |
| Valine | [M+H]+ | 1 | 118.08 | ESI pos | 1.92 | 5.76E-03 | AC vs HP | E |
| Xanthine | [M+H]+ | 2 | 153.03 | ESI pos | 1.39 | 8.90E-10 | AC vs AP, AC vs HP | D |
