## Supplementary figures and images for "Composition of saliva metabolome is significantly associated with SARS-CoV2 infection and with severity of COVID-19 disease"

### Supplemental Figure 1

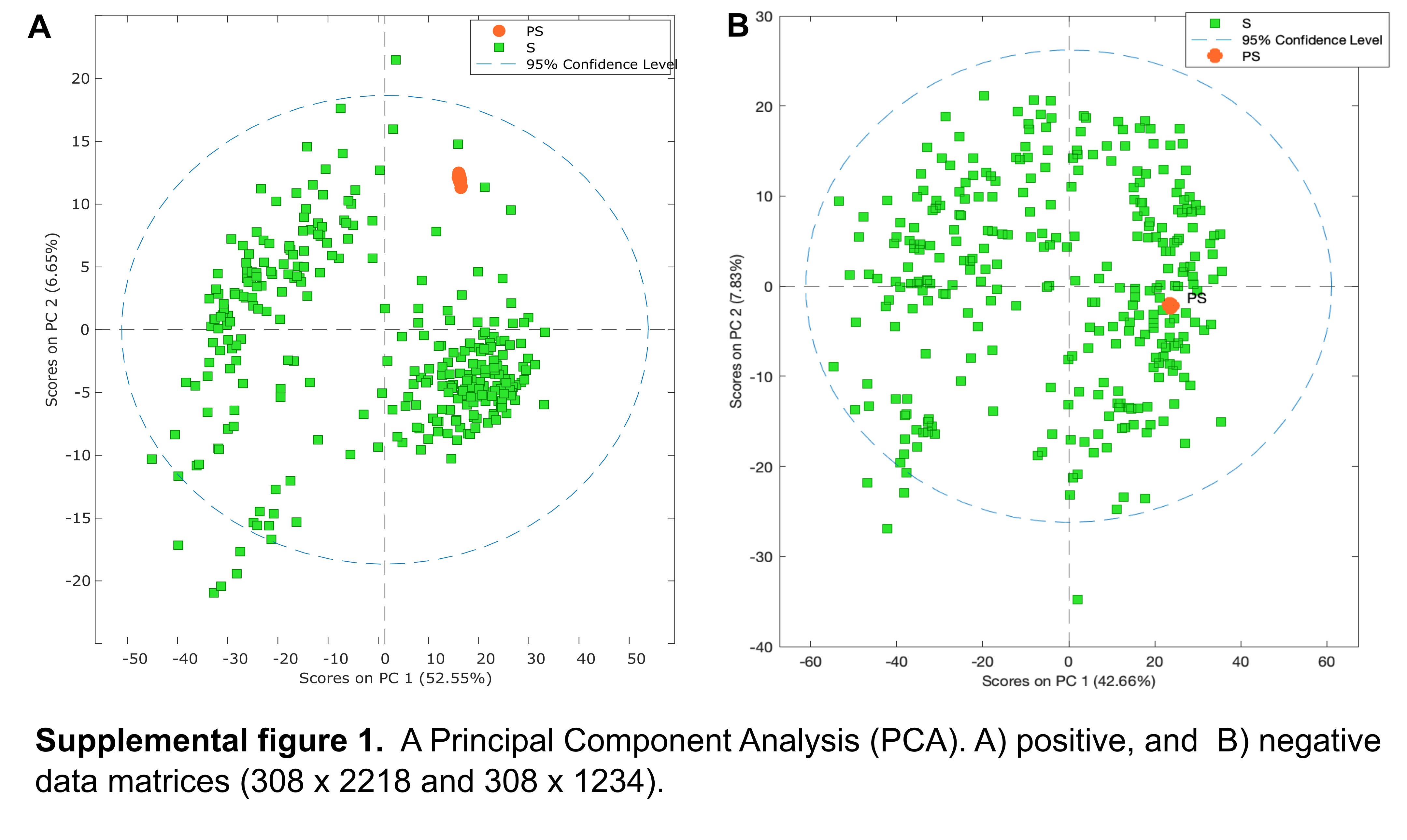

### Supplemental Figure 2

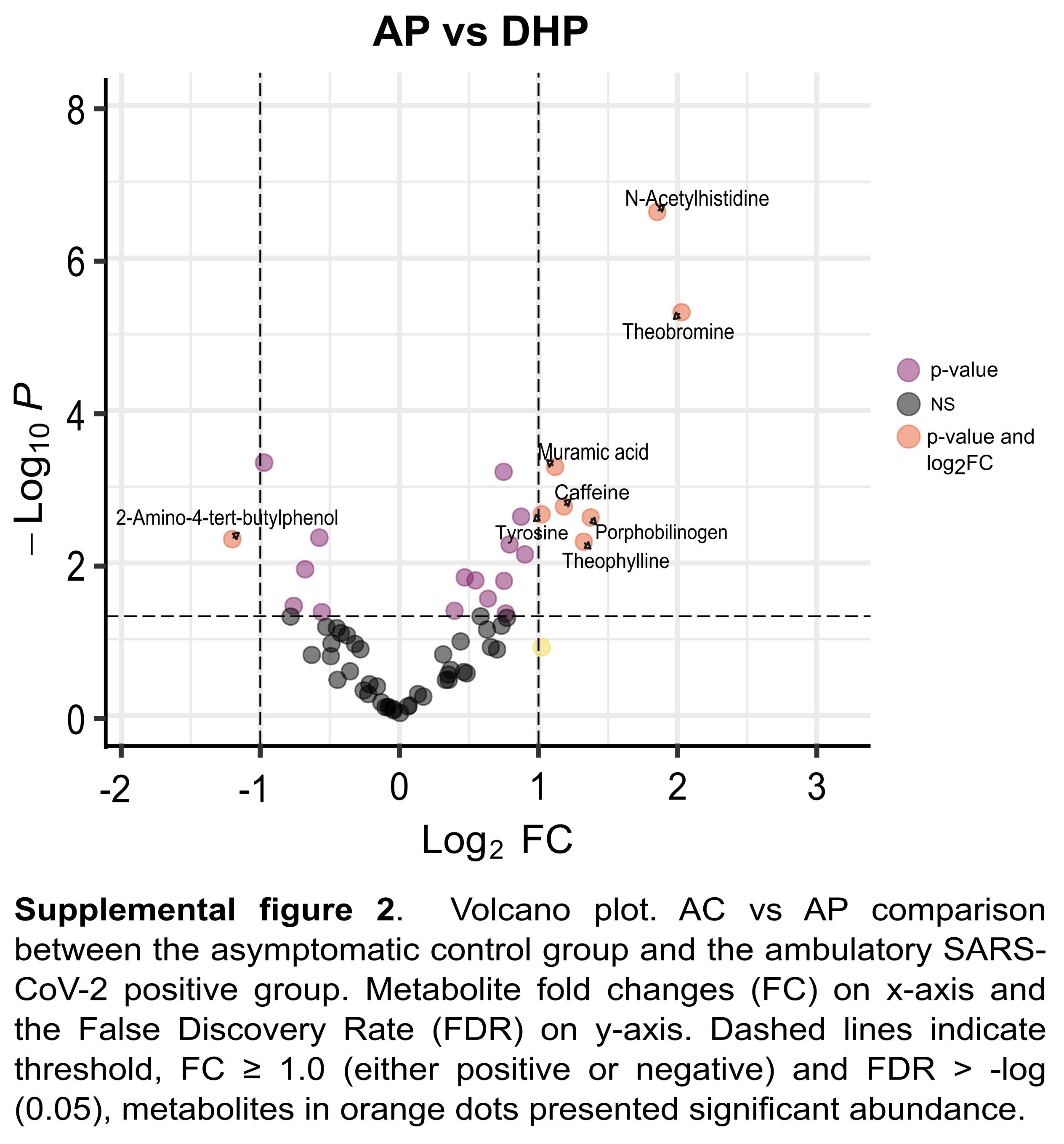
